# RAB1A is a novel vulnerability in uveal melanoma revealed by dual inhibition of MNK1/2 and mTOR

**DOI:** 10.1101/2025.02.23.639782

**Authors:** Raul E. Flores-Gonzalez, Christophe Goncalves, Noah Wirasinghe, Veronique Gaudreault, Jesus M. Noda Ferro, Kelly Coutant, Andrew Mitchell, Elizabeth Guettler, Daniel R. Gonzalez-Moreno, Samuel Preston, Feiyang Cai, Natascha Gagnon, Steve Jean, Solange Landreville, Wilson H. Miller, Sonia V. del Rincón

## Abstract

Uveal melanoma (UM) is an eye cancer that is fatal upon metastasis to the liver. Most treatments trialed in UM fail to provide therapeutic benefit, thus there is an urgent need for novel treatment strategies. The MAPK and PI3K signaling pathways, key molecular drivers found to be hyper-activated in UM, converge on the MNK1/2-eIF4E and mTORC1/2-4EBP axes. Here, we demonstrate that the pharmacologic inhibition of MNK1/2 in combination with an mTOR inhibitor impairs clonogenic outgrowth and UM cell invasion. Using proteomic analyses, we reveal that combined MNK1/2 and mTOR inhibition disrupts Golgi homeostasis and protein vesicle trafficking mainly due to downregulated RAB1A expression, a master regulator of intracellular protein transport. We uncover that the knockdown of RAB1A blocks liver metastasis, a result that is recapitulated by combined pharmacologic inhibition of MNK1/2 and mTOR. Finally, we show that RAB1A expression reshapes the surfaceome by increasing the abundance of plasma membrane proteins associated with poor overall survival in UM, highlighting its potential as a biomarker. This study identifies protein vesicle transport as an unrecognized vulnerability in UM and supports a mechanistic rationale for targeting MNK1/2 and mTOR in metastatic UM.

## INTRODUCTION

Uveal Melanoma (UM) is the most common form of primary intraocular cancer in adults, arising from the melanocytes within the uveal tract. Local management of the disease can be achieved with surgery (enucleation or local resection) or radiation therapy, but more than half of patients diagnosed with UM will develop liver metastasis, at which time their 5-year survival rate is estimated to be about 15% (1). Recent years have seen progress in the treatment of UM, notably with the drug Tebentafusp, with improved overall survival to 21.6 months survival in adults with unresectable or metastatic uveal melanoma, compared to 16.9 months of patients having standard treatment (2). However, only a small subset of patients (HLA-A*02:01 allele with gp100 melanoma marker) may benefit from receiving Tebentafusp. Patients that do not meet the latter criteria, comprising more than 50% of those with UM, have few to no alternative treatment options (3–6). As a result, there is a continuous need for new treatment strategies in UM.

Contrary to cutaneous melanoma, UM rarely harbors mutations in *BRAF* (7); therefore, UM is not treatable with MAPK-targeted therapies as a monotherapy (8). Instead, two mutually exclusive mutations in *GNAQ* and *GNA11*, which encode the G-protein alpha-subunits that mediate signaling downstream from G-protein-coupled receptors, are present in more than 85% of patient tumors (9). Mutations in *GNAQ* and *GNA11* result in the constitutive activation of the PI3K and MAPK signaling pathways, which converge on the eIF4F complex, which controls mRNA translation (10, 11). This complex is comprised of eIF4E (5’-cap binding protein), eIF4G (a scaffolding protein) and eIF4A (a DEAD box RNA helicase) (12). The phosphorylation and availability of eIF4E is tightly controlled. For example, MNK1/2, activated downstream of p38 and ERK MAPK, phosphorylate eIF4E on serine 209, a modification which is critical for the oncogenicity of eIF4E (13). When phosphorylated, eIF4E increases the translation of a subset of mRNAs coding for proteins involved in cell survival and invasion (14, 15). The availability of eIF4E is regulated by binding to 4E binding proteins (4E-BPs) (16). In their unphosphorylated form, 4E-BPs bind eIF4E, thus preventing eIF4E from participating in eIF4F complex formation. The phosphorylation of 4E-BPs by activated mTOR frees eIF4E for binding to eIF4G and participate in mRNA translation(17). Several therapies targeting faulty protein synthesis, including MNK1/2 and mTOR inhibitors, have been or are in clinical trials (18). Interestingly, mTOR inhibitors, as well as other anti-cancer therapies, increase eIF4E phosphorylation, which may limit their anti-tumor benefit (19–24). The activation of the MNK1/2-eIF4E axis promotes the translation of metalloproteases (*MMP3*, *MMP9*), transcriptional regulators (*SNAI1*) and survival signals (*MCL1*, *BCL2*) that will result in increased metastatic and proliferative potential. Thus, mTOR inhibitors may be more effective when combined with MNK1/2 inhibitors, which block eIF4E phosphorylation (22). This concept has been explored in other tumor contexts such as *KRAS*-driven colorectal cancer, chronic myeloid leukemia, and pancreatic cancer (25–27). MNK1/2 inhibitors such as CGP57380, SEL201 and eFT508 have been shown to reduce tumor progression, clonogenic outgrowth and invasive potential of diverse tumor types (28–31).

We hypothesized that UM are driven, at least in part, by dysregulated mRNA translation. Herein, we show that the MNK and mTOR axes are druggable in human UM. The use of combined small molecule inhibitors targeting MNK and mTOR decreased clonogenic outgrowth and invasion of several UM cell lines. To mechanistically understand how the inhibitors of protein synthesis impact UM, we performed data-independent acquisition mass spectrometry (DIA MS), which, coupled with Metascape and STRING pathway analysis, revealed disruption of vesicular protein trafficking as a therapeutic vulnerability. We next showed that the small GTPase RAB1A was a downstream target of dual inhibition of MNK and mTOR. In pre-clinical mouse models of UM, genetic repression or overexpression of RAB1A correlated with the presence of liver metastasis. Analysis of publicly available UM patient data [i.e. TCGA-UVM database(32)], showed that patients whose tumors expressed high levels of RAB1A had decreased overall survival, compared to those patients whose tumors expressed low RAB1A. In addition, Class 2 UM samples, i.e. samples most likely to metastasize (33), have higher RAB1A expression when compared to Class 1 UM (34, 35). We also describe that the combination therapy of MNK and mTOR inhibitors decreases metastasis of injected UM cells in an experimental model of liver metastasis. Finally, we demonstrate that targeting RAB1A expression impacts the secretory pathway and reshapes the surfaceome, which may contribute to reduced invasive metastatic potential. Our work reveals RAB1A as a new vulnerability in UM.

## MATERIALS AND METHODS

### Cell culture

UM cell lines were obtained from Pr. Solange Landreville at Université Laval (Uveal Melanoma Biobank, Vision Sciences Research Network). HEK293FT (Thermo Fisher Scientific, #R70007), T108, T128, T142, and T143 cell lines were cultured in DMEM(36–39) (Wisent, #319-005-CL). 92.1, MEL270, MEL290, MU2, MU2F, OMM2.5 and MP46 were cultured in RPMI 1640(40–42) (Wisent, #350-000-CL). H79 was cultured in DMEM low glucose (43) (Wisent, #319-010 CL). Culture media were supplemented with 10% FBS, 100 IU/mL penicillin and 100 IU/mL streptomycin (Wisent, #450-201-EL) grown at 37°C and 5% CO_2_. During experiments, cells were exposed to 2.5 µM SEL201 (RYVU Therapeutics) and 25 nM INK128 (Cayman Chemical). For subculturing or at endpoint, trypsin 1X (WISENT, #325-242-EL) was added after washing with PBS (WISENT, #311-425-CL) followed by addition of 7 mL of complete media to inactivate the trypsin. Cells were spin for 5 min at 240g and resuspended in 3 to 5 mL of complete medium or PBS for downstream experiments.

### Colony formation assay

Cells were seeded in 6 well plates at 250 to 2,000 cells per well and allowed to adhere overnight. The medium was changed and drugs added the next day. The medium with drugs was changed every 2 days. At the experimental endpoint, cells were stained with 0.5% crystal violet in 70% ethanol, the plates scanned, and colonies were manually counted using the ImageJ software (https://imagej.net/ij/). For each cell line, we performed at least 3 biological replicates, and 3 triplicates per condition.

### Invasion assay

Cells were seeded in a 15 cm petri dish at 1 million cells per plate and allowed to adhere overnight. Cells were starved (serum-free medium) for 16h and then seeded in 12 well plates from 50,000 to 100,000 cells per well in an upper chamber of a polycarbonate insert (8 µm pore size, Corning, # 353182) previously coated with Matrigel (Corning, #354230). At experimental endpoint, the polycarbonate inserts were washed with PBS and cells fixed with 5% glutaraldehyde solution followed by staining with 0.5% crystal violet in 70% ethanol. Non-migratory cells were removed, and invasive cells were counted using EVOS M5000 automatic counting software and manual counting using ImageJ. For each cell line, we performed at least 3 biological replicates, and 3 triplicates per condition. For experiments involving siRNA-mediated knockdown, cells were seeded in 6 well plates at 100,000 cells per well and allowed to adhere overnight. A master mix solution consisting of 3.25 mL of OPTIMEM (Thermo Fisher Scientific, #11058021), 32.5 µL of Lipofectamine RNAiMAX (Invitrogen, #13-778-150) and 6 µL of siRNA targeting *RAB1A* (IDT, siRNA#1 GAUGUCAGGUUUAGUCUUCUGAAGA, siRNA#2 GAUUAUUUAUUCAAGUUACUUCUGA) was prepared, mixed and incubated for 20 min at room temperature. Five hundred uL of the master mix were added to each well along with 2 mL of medium. After 24h, 2 mL of fresh medium were added to each well. Cells were harvested at 48h post-transfection, counted and resuspended in serum-free medium.

### Western blotting

A complete protein lysate buffer was prepared using RIPA containing 10 mM Tris (pH 7.4), 1% NP40, 0.1% SDS, 0.1% sodium deoxycholate and 150 mM sodium chloride. Protease and phosphatase inhibitors were added (Roche, #11697498001 and #4906845001) prior to lysing cell pellets. Then, cells were lysed using 50 to 150 uL of complete RIPA and sonicated at 50% power for 4 seconds before centrifuging at 15 000g for 15 min. The protein concentration was measured by Bradford assay (Biorad, #500-0006). Equal levels of proteins were loaded and separated on 10% SDS-PAGE, then transferred into nitrocellulose membranes, and blocked for 1h in 5% non-fat milk dissolved in TBS-T buffer. Membranes were incubated with primary antibodies overnight at 4°C. The next day, membranes were washed with TBS-T buffer and incubated with a secondary antibody for 1h, followed by 3 washes of 10 min with TBS-T. Membranes were developed using Amersham ECL Prime Western Blotting Detection Reagent (Cytiva, #RPN2106) and Immobilon Western Chemiluminescent HRP Substrate (Millipore, #WBKLS0500).

### Access and reanalysis of previously published data sets

scRNA-seq data were downloaded from the NCBI Gene Expression Omnibus (GEO) public database, accession number GSE139829(35) and GSE138665(34). RAB1A expression in the uveal melanoma cell population of each patient was further examined after normalization.

### RAB1A KD and OE cell lines

HEK293FT cells were seeded in a 10 cm petri dish at 2 million cells per plate and allowed to adhere overnight. Cells were transfected with a packaging plasmid (psPAX2), a viral envelope expression plasmid (MD2.G), and either pLKO Control vector or pLKO-shRNA with constructs targeting *RAB1A* (*Homo sapiens* NM_004161.5, TRCN0000157760 and TRCN0000280677), or pLX317 control vector and pLX317 RAB1A ORF for gene overexpression of *RAB1A* (*Homo Sapien*s, ORF TRCN0000466775). T128 and T143 UM cell lines were seeded in 10 cm dishes at 1 million cells per dish. The next day, viral particles present in the HEK293FT medium were harvested. A total of 4 mL of medium with viral particles was used to tag the cell lines. Cells were treated with 0.5 ug/mL of puromycin (Sigma, P8833-25MG) to select resistant cells, then further expanded.

### Luc-tdTomato labeled cell lines

HEK293FT cells were seeded in a 10 cm petri dish at 2 million cells per plate and allowed to adhere overnight. Cells were transfected with a packaging plasmid (psPAX2), an expression plasmid (pCDH-EF1-Luc2-P2A-tdTomato), and a viral envelope expression plasmid (MD2.G). T128 UM cells (pLKO, pLKO-shRNA, pLX317 and pLX317 RAB1A OE) were seeded in 10 cm dishes at 1 million cells per dish. The next day, viral particles present in the HEK293FT medium were harvested. A total of 4 mL of medium with 8 µg/mL of polybrene (Sigma, TR-1003-G) with viral particles was used to tag the cell lines. Cells tdTomato+ were bulk sorted and expanded.

### TCGA patient survival analysis

Overall survival data from each patient (N=80, divided in low expressing and high expressing tumors based on RNA expression) were obtained from TCGA-UVM (UCSC Xena Browser) using gene expression dataset. Data for the genes *RAB1A*, *CADM1*, *EPHB2* and *LRP8* were graphed using GraphPad Prism KM survival.

### Mass spectrometry sample preparation

T128 cells were seeded in a 15 cm petri dish at 2 million cells per plate and allowed to adhere overnight. Cells were treated with inhibitors and harvested at 24h post-treatment. We performed 4 biological replicates per treatment. Frozen cell pellets were resuspended in buffer (10 mM HEPES pH 8, 8 M urea), and quantified using BCA assay (Thermo Fisher Scientific, #23225). To reduce disulfide bonds, dithiothreitol (DTT) was added to a final concentration of 5 mM in 50 μg of protein, and samples were heated at 95°C for 2 min. Following a 30-min incubation at room temperature, chloroacetamide (7.5 mM) was added for alkylation, and samples were incubated in the dark for 20 min at room temperature. To dilute the urea concentration to 2 M, 50 mM ammonium bicarbonate (NH□HCO□) was added. Peptide digestion was carried out by adding 1 μg of trypsin (Trypsin Gold, Promega) and incubating the samples overnight at 30°C. The reaction was quenched by acidifying the samples to a final concentration of 0.2% trifluoroacetic acid (TFA). An aliquot of the digested peptides was set aside at this stage for DIA spectral library generation. The remaining samples were purified using ZipTip C18 columns (EMD Millipore), lyophilized using a speed vacuum concentrator, and reconstituted in 1% formic acid. The peptide concentration was determined by measuring the absorbance at 205 nm using a NanoDrop spectrophotometer (Thermo Fisher Scientific). Samples were then transferred to glass vials (Thermo Fisher Scientific) and stored at −20°C until mass spectrometry analysis.

### DIA spectral library preparation

Aliquots set aside for the DIA spectral library were desalted as previously described in the previous section, then dried in a speed vacuum concentrator, and resuspended in 300 μL of 0.1% TFA. Fractionation of the peptide mixture was done using the Pierce High pH Reversed-Phase Peptide Fractionation Kit (Thermo Fisher Scientific). Each fractionation column was first centrifuged at 5,000 × g for 2 min at room temperature to remove excess liquid and to pack the resin. The column was then washed twice with around 300 μL of 100% acetonitrile (ACN) and conditioned with two additional washes of 0.1% TFA. Purified peptides were loaded onto the column, then centrifuged at 3,000 × g for 2 min at room temperature and washed with 300 μL of mass spectrometry-grade water. Peptides were sequentially eluted in eight fractions by applying 300 μL of solutions containing 0.1% triethylamine and increasing ACN concentrations (5% to 50%). Each elution step was followed by centrifugation at 3,000 × g for 2 min at room temperature, with fractions collected in separate low-binding microtubes. Fractions were then concentrated to dryness using a speed vacuum concentrator at 60°C and resuspended in 50 μL of 1% formic acid (FA). Peptide concentrations were determined by measuring the absorbance at 205 nm using a NanoDrop spectrophotometer (Thermo Fisher Scientific). Finally, peptides were transferred to glass vials (Thermo Fisher Scientific) and stored at −20°C until mass spectrometry analysis.

### DIA LC-MS analysis

For each sample, 250 ng of peptides were injected into the nanoElute high-performance liquid chromatography (HPLC) system (Bruker Daltonics), and later, loaded onto a trap column (Acclaim PepMap100 C18, 0.3 mm ID × 5 mm, Dionex Corporation) with a constant flow rate of 4 μL/min. Peptides were then eluted onto an analytical C18 column (1.9 μm bead size, 75 μm × 25 cm, PepSep) using a 2h gradient of ACN from 5% to 37% in 0.1% FA at a flow rate of 400 nL/min. Eluted peptides were analyzed using a TimsTOF Pro ion mobility mass spectrometer (Bruker Daltonics) equipped with a Captive Spray nano-electrospray ionization (ESI) source. The data acquisition was performed in data-independent acquisition (DIA) mode using dia-PASEF. Each ion mobility separation cycle (100 ms per trapped ion mobility spectrometry, TIMS, event) was acquired in dia-PASEF mode, with a single mobility window consisting of 27 mass steps per cycle (1.27 s duty cycle). These steps spanned a mass-to-charge ratio (m/z) range of 114 to 1,414 with a mass width of 50 Da, covering the diagonal scan line for doubly and triply charged peptides in the m/z-ion mobility plane. The final target intensity was set to 20,000 and intensity threshold of 2,500.

### Protein identification using MaxQuant analysis with TIMS MaxDIA

Mass spectrometry data were analyzed using MaxQuant (version 2.0.3.0) with the UniProt human proteome database (March 2020 release, 75,776 entries). The TIMS MaxDIA workflow was applied with all raw files assigned to the same cell type and fractionated from 1 to 8. Trypsin was specified as the digestion enzyme (cleaving after K/R except when followed by P), allowing for one missed cleavage. Carbamidomethylation of cysteine was set as a fixed modification, while methionine oxidation and N-terminal protein modifications were included as variable modifications. A mass tolerance of 20 ppm was used for both precursor and fragment ions. To ensure high-confidence identifications, peptide-spectrum matches (PSM FDR), protein identifications (Protein FDR), and site decoy fractions were filtered using a 0.05 false discovery rate (FDR) threshold. A minimum of one peptide was required for protein identification. Additionally, the “Second peptides” and “Match between runs” options were enabled. The analysis was conducted in library mode with a transfer q-value threshold of 0.3, incorporating the "peptides.txt," "evidence.txt," and "msms.txt" files generated from the spectral library. For quantification, label-free quantification (LFQ) normalization was applied across all samples. After identification and normalization, proteins flagged as "Reverse," "Only identified by site," or "Potential contaminant" were excluded. Proteins detected by a single unique peptide were also removed from further analysis.

### MS data analysis for LFQ-MS

LFQ relative intensities obtained from MaxQuant were processed for statistical analysis. PERSEUS software was used to perform multiple hypothesis testing using two-sided Student’s t-test and permutation-based FDR correction of 10%. Significant up/downregulated proteins were normalized and hierarchically clustered. Pathway analyses were performed for each cluster using Metascape to determine the effect of the drug combination on specific pathways.

### Splenic injection model

Six-week-old NCG mice (Charles River Laboratories) were injected in the spleen with 100,000 T143 or T128 luc-tdTomato cells resuspended in PBS to induce metastasis in the liver. Splenectomy was performed only in cohorts treated with vehicle or SEL201 and INK128. Mice were imaged using AMI HT (Spectral Imaging) by injecting 200 uL of luciferin (15 mg/mL) (PerkinElmer, #122799) dissolved in sterile PBS weekly or bi-weekly to tract metastatic growth. At the endpoint, mice were euthanized, the liver and lungs were harvested and fixed in 10% formalin before paraffin embedding for IHC assays. Surface liver metastases were visually quantified for T143 mice cohorts the day after euthanasia.

### TMA IHC and scoring

Cores from human primary UM FFPE tissues were acquired from 18 patients and arranged/mounted in a TMA. Only 11 patient cores had sufficient tissue to be analyzed. Four-μm TMA sections were stained for p-eIF4E, p-S6, and RAB1A using IHC (see below). Cores were scored using the following metric: low, medium, or high expression, based on the overall cell intensity signal quantified using the QuPath software (https://qupath.github.io/). For p-eif4E, a low score corresponded to an intensity range of 0 to 0.1, medium score ranged from 0.1 to 0.16, and high score was above 0.16. For p-S6, low score ranged from 0 to 0.125, medium score from 0.125 to 0.22, and high score was above 0.22. For RAB1A, low score ranged from 0 to 0.12, medium score from 0.12 to 0.23, and high score was above 0.23. Correlation was determined using a Spearman’s test. Red fluorescent protein (RFP) and RAB1A stainings were performed in murine liver tissues. Metastatic UM cells expressing tdTomato were quantified using QuPath. The metastasis area was calculated by dividing area covered by metastases over total liver tissue area.

### IHC on livers bearing UM metastases

NCG mouse livers were collected, washed with 1X PBS, and fixed in 10% neutral buffered formalin (VWR, # 89370-094). Samples were then embedded in paraffin, sectioned at a thickness of 4 μm, and mounted on Superfrost slides. IHC staining was conducted on liver tissues containing metastases. Slides underwent deparaffinization, rehydration, and heat-induced antigen retrieval. To inhibit endogenous peroxidase activity, the tissues were treated with 10% hydrogen peroxide for 15 min, followed by blocking with 10% donkey serum (Jackson ImmunoResearch, #017-000-121). The primary antibody staining was performed using nucleolin (metastases; 1:2,000 dilution), p-eIF4E (metastasis-bearing livers; 1:50 dilution), p-S6 (metastasis-bearing livers; 1:200 dilution), and RAB1A (1:100 dilution). The Magenta substrate (Agilent, #GV92511-2) or the ImmPACT DAB substrate (Vector Laboratories, #SK-4105) was used for visualization with Harris-modified hematoxylin (EMD Millipore, #638A-85) as the counterstain. Images of the stained slides were captured using a Zeiss AxioScan.Z1 slide scanner and analyzed with the QuPath software.

### Retention Using Selective Hooks (RUSH) assay

T128 cells were cultured in DMEM supplemented with 10% FBS and pen/strep with 2ug/ml of puromycin. Cells were detached with trypsin + 0.25% EDTA and a total of 3×10^4^ cells were plated on round glass coverslips (72230-01; Electron Microscopy Sciences) in a 24-well plate. Cells were allowed to adhere for 48 h before being transfected with 500 ng of Ii-Str_SBP-EGFP-GPI plasmid (65296, Addgene) using JetPRIME reagent (101000046; Polyplus), as per the manufacturer’s instructions. The transfection media was removed 7 h later and replaced with Opti-Minimal Essential Medium (MEM) supplemented with 2% FBS. Twenty-four hours after the transfection and 1h before the RUSH assay, the culture medium was replaced with DMEM supplemented with 10% of dialyzed FBS, to limit the amount of biotin in the medium. The RUSH assay was initiated by replacing the culture medium with DMEM supplemented with 10% dialyzed FBS and 40 µM of biotin. Cells were incubated at 37°C for 0, 10, 20 or 30 min. Plates were then lifted on ice and medium was quickly replaced by ice-cold PBS. Cells were fixed for 15 min at room temperature with 4% paraformaldehyde (Electron Microscopy Science, #15713) in PBS, and then incubated for 1h at room temperature in PBS containing 0.3% Triton X-100 and 1% bovine serum albumin (BSA). Two sets of cells incubated without the addition of biotin during the RUSH assay (time 0) were labelled to identify different proteins. The first set was stained with anti-HA (1:800; Cell Signaling, #3724), anti-GFP (1:1,000; Santa Cruz Biotechnology, #sc-9996) and DAPI to evaluate the colocalization of the cargo and hook. The second set was stained with the rest of the RUSH assay samples with anti-GM130 (1:3,000; Cell Signaling, #12480), anti-GFP and DAPI to evaluate the colocalization of the cargo and cis-Golgi. Secondary antibodies used were Alexa-Fluor 488-conjugated anti-mouse and Alexa-Fluor 546-conjugated anti-rabbit (1:400; Invitrogen, #A11029 and #A11035). Cells were mounted on slides with 3 µL of slow fade gold mounting medium containing DAPI (Thermo Fischer Scientific, #S36938). Slides were kept at 4°C in the dark before image acquisitions on a Zeiss LSM880 equipped with a 40x 1.4 plan Apo objective. Images were exported as tiff files using Zen Blue, and then processed using Adobe Photoshop. All images were thresholded and cropped similarly in Photoshop for the experimental set. Figures were assembled using Adobe Illustrator. The image analysis was performed using Cell Profiler (https://cellprofiler.org/). Pearson coefficients were calculated per cell using the colocalization module. All experiments were performed three times independently. The colocalization of the cargo and hook at time 0 was statistically compared between cell types using a Kruskal-Wallis test followed by Dunn’s multiple comparison test. The colocalization of the cargo and Golgi at the 4 tested time points was analyzed by a two-way ANOVA followed by Dunnett’s post-test.

### Surface labeling

Solutions and cell plates were prechilled at 4°C, and all steps were performed on ice to prevent the internalization of membrane proteins. Cells were washed twice with Buffer A (0.5 mM MgCl□, 1 mM CaCl□ in 1X PBS, pH 7.4) and then incubated for 30 min at 4°C in the dark with 1 mM sodium metaperiodate (Sigma-Aldrich, #S1878-25G) dissolved in Buffer A. Subsequently, cells were washed twice with cold Buffer A and incubated for 1h at 4°C in the dark with 100 μM aminooxy-biotin (Biotium Inc., #90113) and 10 mM aniline (Sigma-Aldrich, #242284) for biotinylation. Cells were washed twice again with Buffer A and gently scraped off the plate surface with lysis buffer containing a protease inhibitor cocktail (Sigma-Aldrich, #P8340). The lysate was gently rocked at 4°C for 30 min, and vortexed every 5 min. Nuclei and cell debris were removed by centrifugation at 2,800 × g for 15 min, followed by 16,000 × g for 15 min. The supernatant was collected for membrane protein isolation with high-capacity streptavidin resin (Thermo Fisher Scientific, #20357).

### Sample preparation and mass spectrometry analysis

After several harsh washes (see complementary protocol), streptavidin beads were washed five times with 20 mM ammonium bicarbonate. Proteins bound to beads were reduced with 10 mM DTT in 20 mM NH_4_HCO_3_ and alkylated with 15 mM chloroacetamide in 20 mM NH_4_HCO_3_. The digestion was carried on overnight at 37°C with 1 µg of trypsin (Thermo Fisher Scientific, #PI90058). Beads were finally centrifugated and the supernatant containing the peptides was collected. The peptide mixture was dried and resuspended in 300 μL of 0.1% TFA to subsequent desalting on ZipTip micropipette tips containing a C18 resin (EMD Millipore). The MS analysis was performed by diaPASEF technique implemented in the timsTOF from Bruker Daltonics. For each sample, 250 ng of the desalted peptide mixtures were analyzed by a LC-MS system (nanoElute, Bruker Daltonics), with diaPASEF on a timsTOF Pro ion mobility mass spectrometer. Peptides were loaded onto a trap column (Acclaim PepMap100 C18 column, 0.3 mm inner diameter × 5 mm, Dionex Corporation) at a flow rate of 4 μL/min and moved to an analytical C18 column (1.9 μm particle size, 75 μm x 25 cm; PepSep) maintained at 50°C. Then, peptides were eluted over 120 min gradient time (ACN (5%–37%) in 0.1% FA at 400 nL/min) and injected into the Captive Spray nano electrospray source of TimsTOF Pro (Bruker Daltonics). In diaPASEF mode, a mobility window consisting of 27 mass steps (m/z between 114 to 1,414 with a mass width of 50 Da) per cycle (1.27 seconds duty cycle) was used. Peptides carrying +2 and +3 charges were selected for analysis.

### MS data analysis

Protein identification was performed with DIA-NN (version 1.9.1). *In silico* spectral libraries were created using the Human FASTA file (one protein per gene) from the UniProt database, with Proteome ID UP000005640 and the universal protein contaminant FASTA file available on the repository (https://github.com/HaoGroup-ProtContLib; Frankenfield *et al.*, 2022). Deep learning-based spectra, RTs and IMs prediction option, as well as N-terminal M excision, C(cam) and M(ox) modifications were enabled. Moreover, trypsin/P was selected allowing two missed cleavages and the precursor charge range was set to 2+ and 3+. Bruker.d raw data were analyzed by searching against the generated libraries.

### Differential expression analysis

Membrane proteins were selected by their accession numbers on UniProt. A total of 846 membrane proteins were identified with DIA-NN. Differential expression analysis (DEA) between biological conditions was achieved using the open-source R package Mass Spectrometry Downstream Analysis Pipeline (MS-DAP). This package allows the analysis of label-free proteomics datasets from the DIA-NN tool. MS-DAP incorporates statistical models for DEA like MSqRob (Goeminne *et al.*, 2016) and MS-EmpiRe (Ammar *et al.*, 2019) that work directly at the peptide level. These novel statistical models are more sensitive in detecting small variations (20-25% change) in peptide abundances, rather than averaging all values into a single protein estimate (as Perseus).

### Defining statistically significant hits

Peptide-level-based models increase sensitivity for detecting differentially expressed proteins but may also inflate false positives. To address this, DEA results were initially restricted to proteins identified with at least two peptides. Furthermore, instead of applying an arbitrary log2 fold change threshold, statistically significant hits were defined automatically through bootstrapping analyses using MS-DAP. This combination of restrictions has been shown to effectively control false positive rates. Finally, all differentially expressed proteins were filtered to retain only proteins with a q-value < 0.01 (Benjamini-Hochberg method).

## RESULTS

### The MNK1/2 and mTORC1/2 kinases are active in human UM

MNK1/2 (hitherto MNK) and mTOR are critical regulators of eIF4E phosphorylation and availability, respectively (44). To determine the activation status of these pathways in primary human UM, we performed immunohistochemistry (IHC) staining on patient-derived UMs, which were arrayed on a tumor microarray (TMA). Using antibodies to detect p-eIF4E and p-S6, downstream targets of MNK and mTOR, respectively, our analysis revealed that most of the UM cores co-expressed p-eIF4E and p-S6 (Fig. 1A, B and Supplementary Fig. S1A). In addition, we show that tumors expressing high p-eIF4E and p-S6 were from patients who had metastases (Fig. 1C). These findings suggest that activated MNK and mTOR signaling may play a role in metastatic disease. Thus, targeting both pathways may provide a therapeutic benefit in a tumor type for which existing therapies are sorely lacking.

**Fig. 1.**
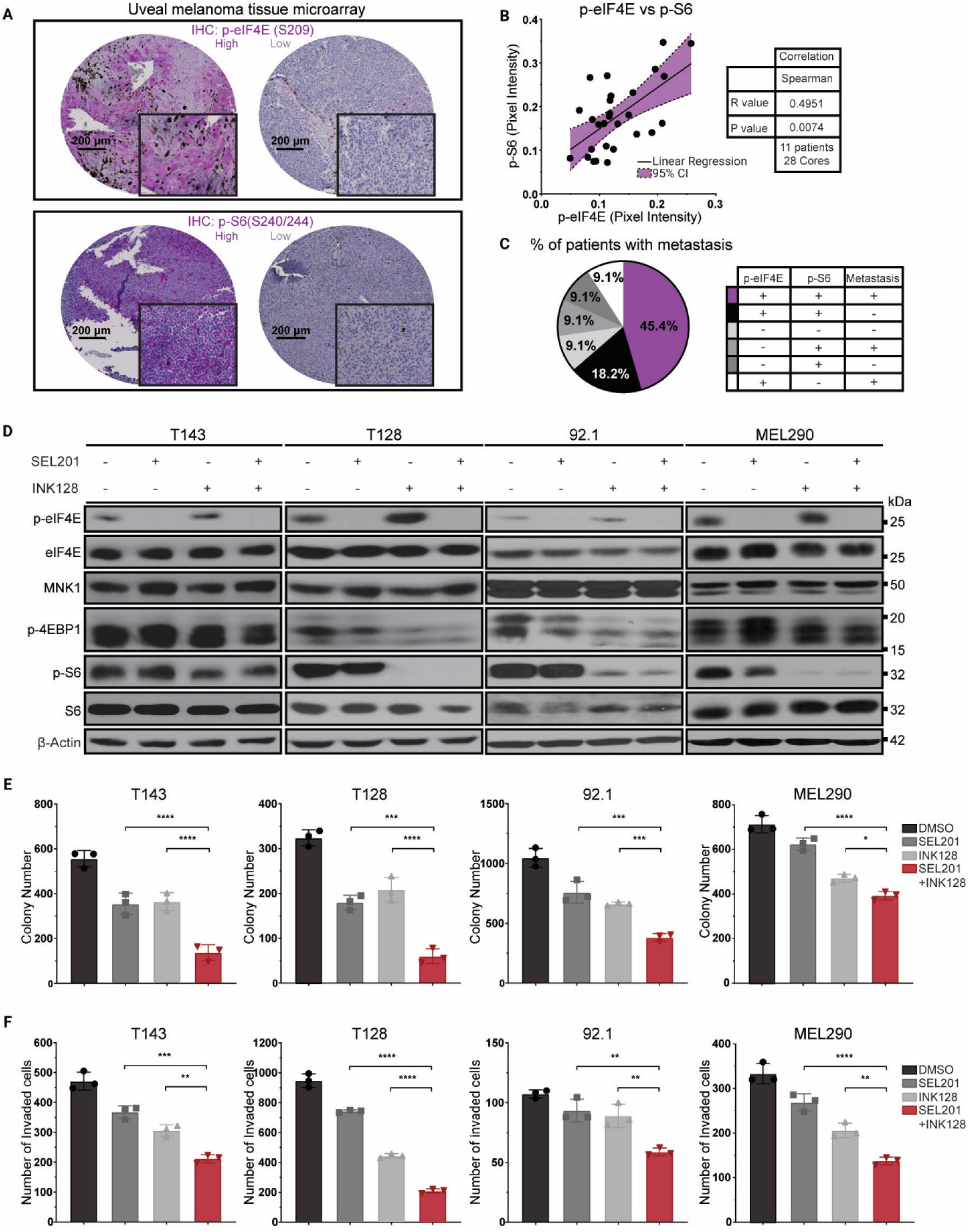
MNK1/2 and mTORC1/2 signaling is active in UM, where its co-targeting decreases clonogenic and invasive outgrowth *in vitro*. **A** Representative images of immunohistochemistry (IHC) assays to detect p-eIF4E and p-S6 signaling status in human primary UM. Linear regression and 95% confidence interval shown in purple. **B** Spearman correlation performed on p-eIF4E and p-S6 tumor cores. **C** Proportion of patients (n=11) that developed metastases (+, -) categorized by p-eIF4E and p-S6 levels (+ high, - low). **D** Immunoblot analysis of the indicated proteins in T143, T128, 92.1 and MEL290 UM cell lines using 2.5µM SEL201 and 25nM INK128 inhibitors, alone or in combination. **E, F** 2.5µM SEL201 cooperates with 25nM INK128 to decrease UM clonogenic outgrowth over 10 days and invasion over 24 hours of UM cell lines. Experiments represent the mean ±SD of 3 experimental replicates performed in technical triplicate. One-way ANOVA multiple comparisons test. \**P* ≤ 0.05, \*\**P* ≤ 0.01, \*\*\**P* ≤ 0.001 and \*\*\*\**P* ≤0.0001 compared to single treatments.

### Pharmacological inhibition of MNK and mTORC1/2 suppresses clonogenicity and invasion

There is some evidence for the anti-tumor benefit of mTOR inhibitors in preclinical UM models (45), but this did not translate into the clinic setting, where resistance is often observed (45). One explanation for the limited clinical benefit of mTOR inhibitors such as Sapanisertib (aka INK128) (46), is the increase in the phosphorylation of eIF4E via compensatory feedback mechanisms (19–22). Accordingly, we hypothesized that therapeutic targeting of mTOR would be more effective when combined with inhibitors of MNK, which block the phosphorylation of eIF4E (22, 47). As expected, INK128 blocked mTOR activity, shown by decreased phosphorylation levels of p-4EBP1 and p-S6. Consistent with previous results, mTOR inhibition led to an increase in eIF4E phosphorylation in some cell lines; however, this effect was mitigated by co-treatment with the MNK inhibitor SEL201 (Fig. 1D). We next tested whether the combined inhibition of both axes alters tumor phenotypes in UM. The combination therapy of MNKi+mTORi decreased invasion and clonogenic potential of UM cell lines, as compared to single agent treatments after 24h and 10 days of treatment, respectively (Fig. 1E, F). Comparable results were observed in seven additional human UM cell lines (Supplementary Fig. S2A-C and Supplementary Fig. S3A-C), showing the broad therapeutic benefit of inhibitors of translational control in UM.

### Data-independent acquisition mass spectrometry (DIA-MS) uncovered pathways regulated in UM cells treated with combined MNK and mTOR inhibitors

DIA-MS, a quantitative proteomic technique, has been widely used to uncover the mechanisms underlying therapeutic action (48). DIA-MS provides increased sensitivity, reproducibility and protein coverage, compared with other quantitative LC/MS-MS approaches (49). We used this proteomic approach to provide insight into the mechanism through which combined MNK and mTOR inhibition represses UM invasion and clonogenic outgrowth. We discovered 326 differentially regulated proteins after 24h of treatment using hierarchical clustering (Fig. 2A). Next, we performed pathway enrichment analysis on each cluster (Supplementary Fig. S4A-F) while focusing on the proteins being downregulated in the combination treatment (Fig. 2B). Particularly, our analysis identified three enriched pathways, two of which are shown to be directly involved in major control of Golgi vesicle trafficking processes comprising membrane organization, vesicle transport and organization (Fig. 2C). Among the proteins that are characteristic of these pathways, we noted an overrepresentation of Rab small GTPase family members, such as RAB1A, RAB8A and RAB13 (Fig. 2D). Of these, RAB1A was chosen for downstream validation studies, as it was previously shown to play a role in cell migration/invasion in other tumor types, but not in UM (50–53). Immunoblotting confirmed that RAB1A protein expression was repressed by combined MNK/mTOR inhibitor therapy in T128, T143, 92.1 and MEL290 UM cells after 24 h of treatment (Fig. 2E). These cell lines were among those in which we observed robust inhibition of clonogenicity and invasion after combined treatment with MNK1/2 and mTORC1/2 inhibitors (Fig. 1E, F).

**Fig. 2.**
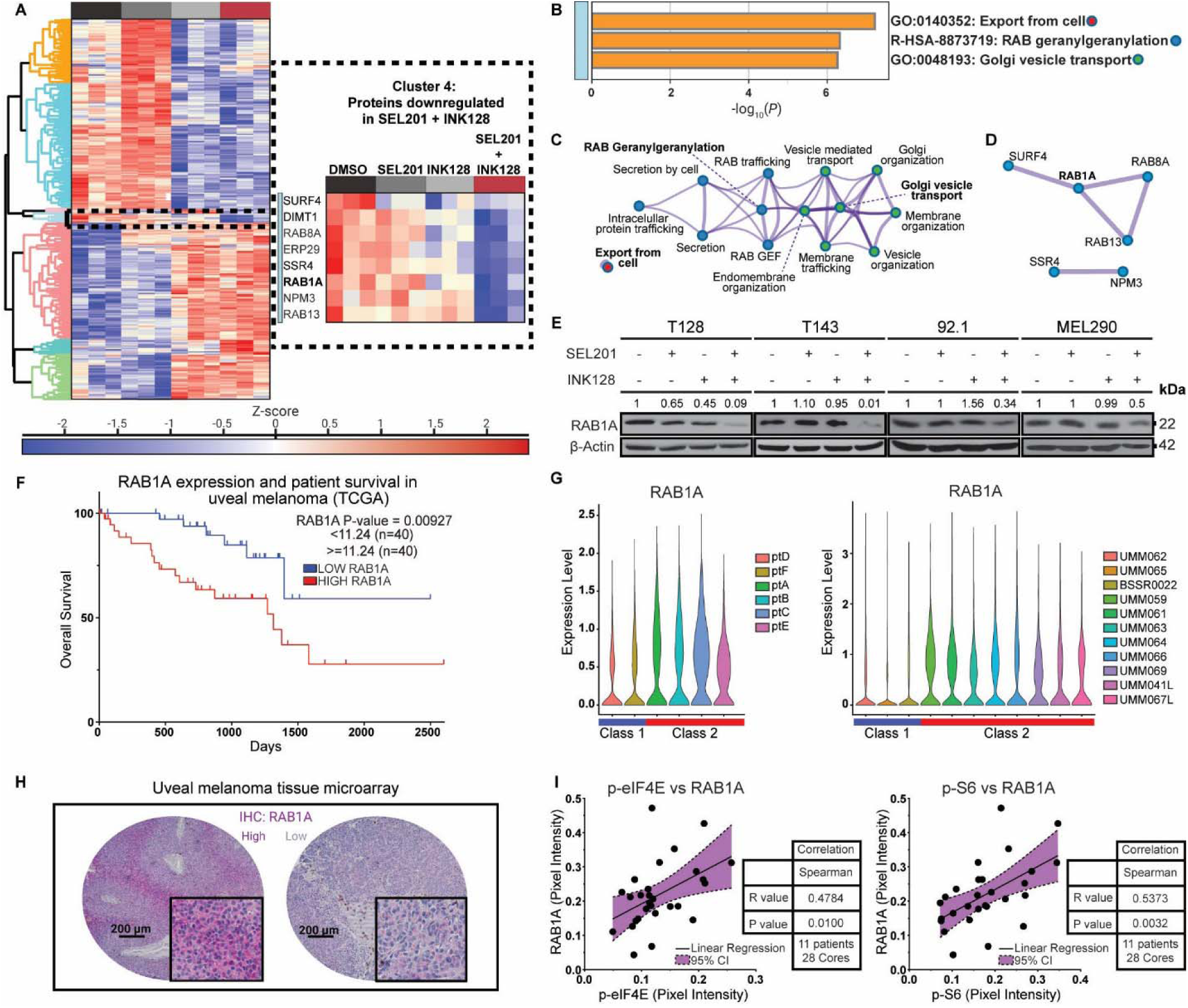
Proteome profiling of T128 UM cell line shows that combined inhibition of MNK1/2 and mTORC1/2 downregulates intracellular protein vesicle trafficking in UM. **A** Heatmap of hierarchical clustering proteome of UM cells treated with vehicle, SEL201, INK128 or combination therapy at 24 h after treatment. **B-D** Pathway Enrichment Analysis and protein-protein interaction of cluster 4 reveals the suppression of key proteins related to Golgi vesicle trafficking processes. **E** Immunoblot validating that combined inhibition of the MNK-eIF4E and mTORC1/2-4EBP1 axes downregulates the expression of RAB1A in T143, T128, 92.1 and MEL290 UM cell lines at 24 hours after treatment. Densitometry values of RAB1A expression relative to DMSO are shown, normalized to β-actin. **F** Kaplan-Meier overall survival plot depicting RAB1A mRNA expression in the TCGA UVM dataset (n=80), *P*-value=0.0092. **G** Single cell RNA expression data from GEO #GSE138665 (left plot) and GEO #GSE139829 (right plot) shows RAB1A expression in Class 1 and 2 UM samples. **H** Representative images of immunohistochemistry to detect RAB1 expression in human UM. **I** Spearman correlation performed on RAB1A vs p-eIF4E (left) and RAB1A vs p-S6 tumor cores (right). Linear regression and 95% confidence interval shown in purple.

### RAB1A is expressed in human UM and correlates with p-S6 and p-4E expression

As the role for RAB1A in UM has not yet been explored, we next interrogated the expression of RAB1A in the TCGA UVM dataset(32), which revealed that increased RAB1A abundance is associated with decreased UM overall survival (Fig. 2F). Moreover, our analysis of two publicly available human UM single-cell RNA sequencing data (34, 35) showed that Class 2 UM, which have a high propensity to metastasize(54), have the highest expression of RAB1A expression when compared to Class 1 UM (Fig. 2G). We next stained the UM TMA described in Fig. 1A for RAB1A using IHC and found that RAB1A expression correlated with p-eIF4E and p-S6 expression (Fig. 2H, I). Together, these data suggest that RAB1A is expressed in human UM, thereby revealing a novel potential therapeutic vulnerability to block metastasis.

### RAB1A expression promotes UM invasion and metastasis

To assess whether RAB1A regulates clonogenic outgrowth and invasion, as observed in UM cells treated with combined MNK and mTOR inhibitors, we engineered UM cells to overexpress or silence RAB1A (Fig. 3A, C, D). Silencing RAB1A using two approaches, siRNA or shRNA (i.e. RAB1A KD), in T128 and T143, was sufficient to inhibit invasion, similar to the results obtained with the drug combination therapy (Fig. 3B, E). In addition, shRNA-mediated knockdown of RAB1A decreased clonogenicity of T128 and T143 cells (Fig. 3F). Conversely, RAB1A overexpression in T128 and T143 resulted in increased cell invasion and clonogenic outgrowth (Fig. 3G, H). We next assessed whether modulation of RAB1A in UM cells would impact liver metastasis, which occurs in more than 90% of patients with UM (1). We injected empty vector (EV) and RAB1A overexpressing (OE) T128 Luc-tdTomato cells into the spleen of NCG mice, a routinely used model of experimental liver metastasis(55) (Fig. 4A). *In vivo* and *ex vivo* imaging revealed higher luminescence, an indicator of greater metastasis burden, in the OE cohort compared to the EV (Fig. 4B-E). We confirmed the greater metastasis burden in the mice injected with OE compared to the EV cells by staining the livers with anti-RFP and human nucleolin IHC staining of livers (Fig. 4F). We further verified the effect of RAB1A in promoting and suppressing experimental liver metastasis using an additional model: T143 UM cells engineered to silence or overexpress RAB1A, respectively (Fig. 4G, H). Our findings reveal that the levels of RAB1A correlate strongly with the presence of liver metastasis.

**Fig. 3.**
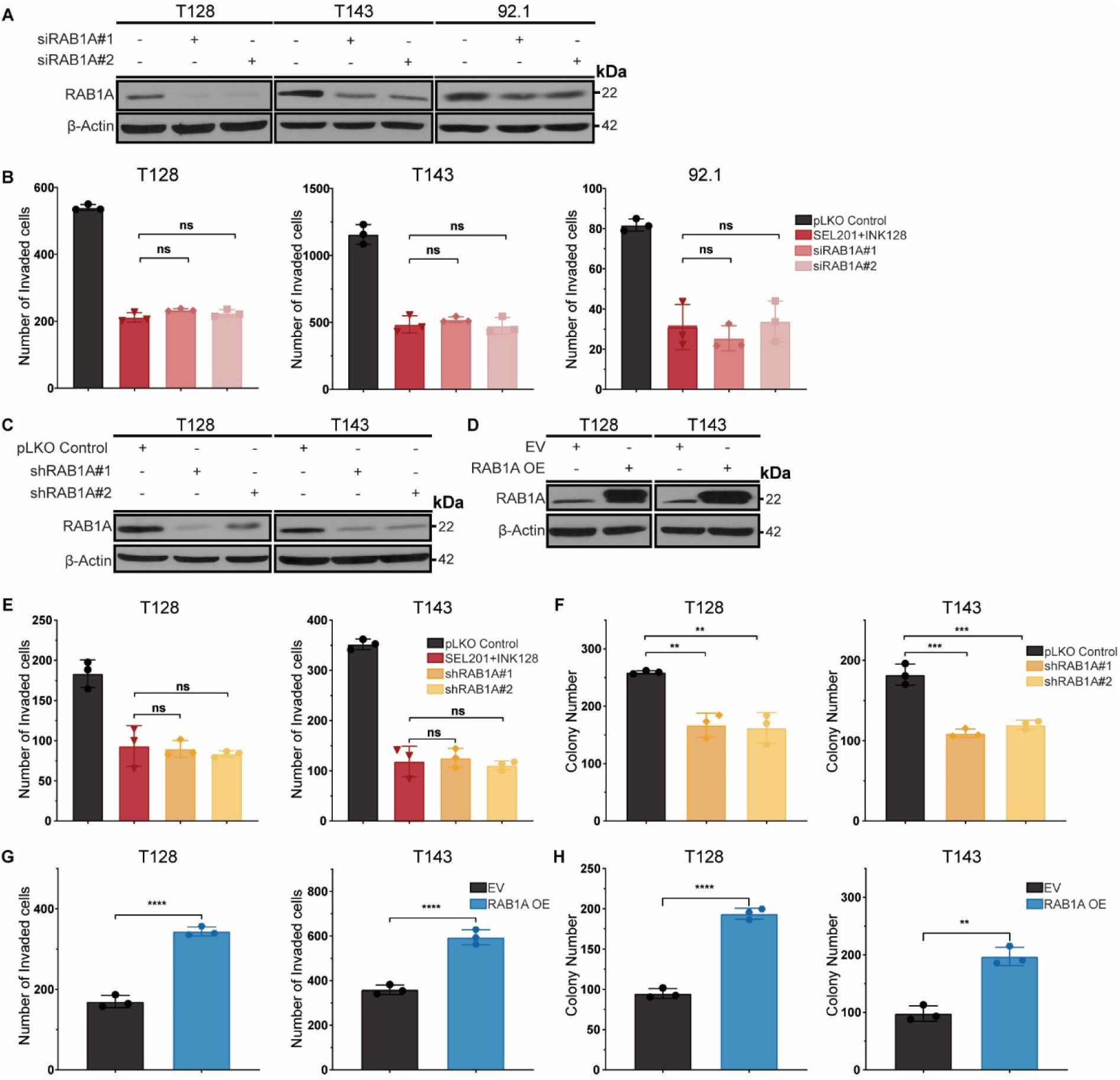
RAB1A expression influences invasive and clonogenic potential of T128 and T143 UM cell lines. **A** Immunoblot of T128, T143, and 92.1 transfected with control treatment or transient siRNA against RAB1A. **B** RAB1A transient knockdown decreases invasion in T128, T143 and 92.1, comparable to the effect observed under combined MNK1/2-mTORC1/2 pharmacological inhibition. Experiments represent the mean ±SD of 3 experimental replicates performed in technical triplicate. One-way ANOVA multiple comparisons test, ns=non-significant compared to combination treatment. **C** Immunoblot of T128 and T143 transduced with stable shRNA (shRNA#1&2) against RAB1A compared to pLKO Control. **D** Immunoblot of RAB1A expression in T128 and T143 cell lines transfected with empty vector (EV) or plasmid encoding human RAB1A (RAB1A OE). **E, F** RAB1A stable knockdown mimics MNK1/2-mTORC1/2 pharmacological inhibition to decrease invasion and clonogenic outgrowth on T128 and T143 cell lines. Experiments represent the mean ±SD of 3 experimental replicates performed in technical triplicate. One-way ANOVA multiple comparisons test, ns=non-significant, \**P* ≤0.05, \*\**P* ≤0.01, \*\*\**P* ≤0.001, \*\*\*\**P* ≤0.0001 compared to combination. **G, H** RAB1A overexpression enhances clonogenic and invasive potential of T128 and T143 compared to EV. Experiments represent the mean ±SD of 3 experimental replicates performed in technical triplicate. Unpaired t-test, \*\**P* ≤0.01, \*\*\*\**P* ≤0.0001 compared to EV.

**Fig. 4.**
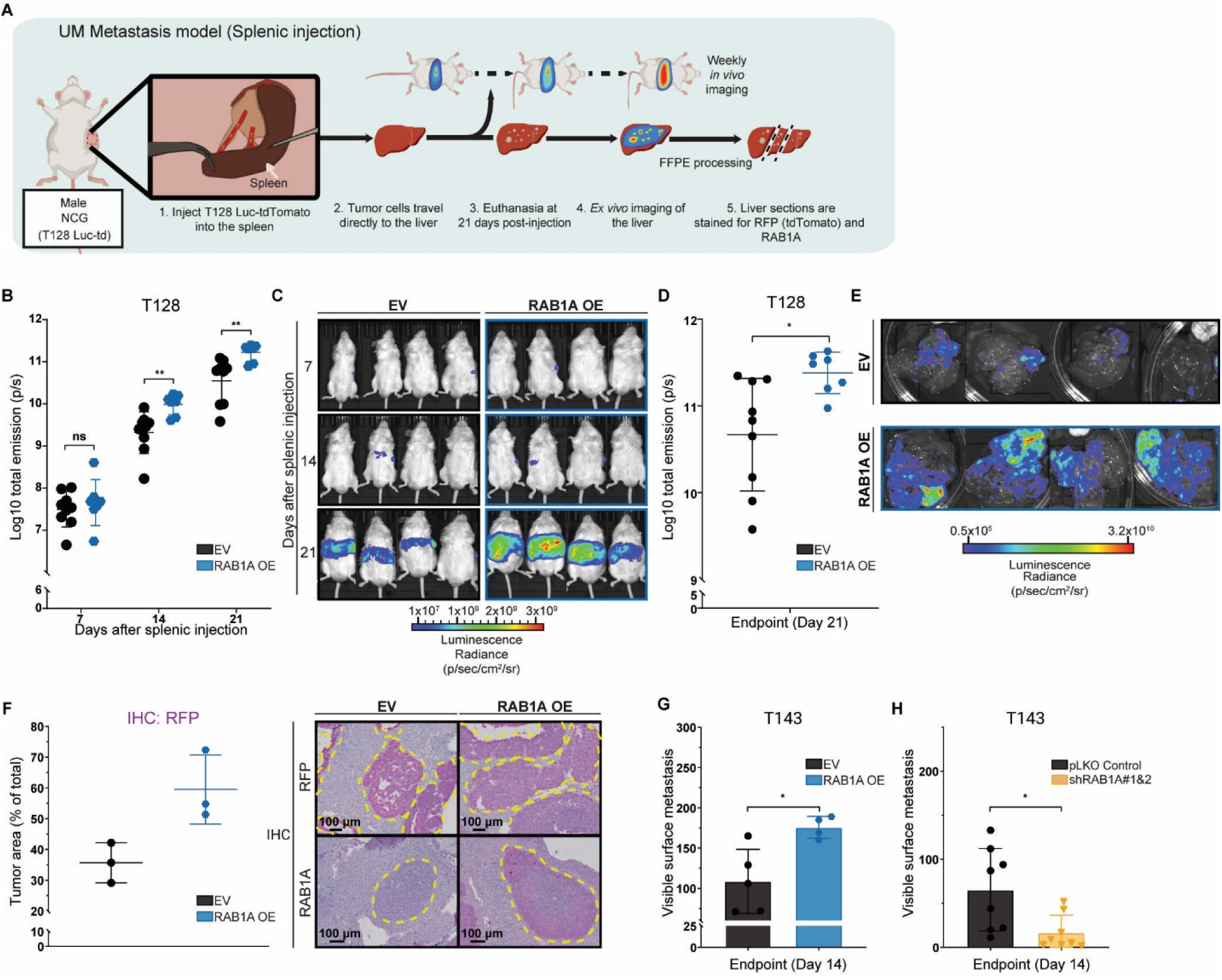
RAB1A expression modulates *in vivo* metastasis potential of T128 and T143 UM cell lines. **A** Splenic injection model diagram using NCG mice injected with T128 luc-tdTomato or T143. Tumor burden (T128) was determined by log10 luminoscore (photons/sec) acquired weekly and using surface metastasis (T143) on NCG livers. **B, C** *In vivo* imaging and representative images at each timepoint of NCG bearing T128 luc-tdTomato tumors transduced with EV (n=9) and RAB1A OE (n=7), Multiple t-test analysis. \*\**P* ≤0.005 at day 14 and 21. **D, E** *Ex vivo* imaging and representative images of NCG livers with T128 luc-tdTomato EV or RAB1A OE tumors, Mann-Whitney test analysis. \**P* ≤0.05. **F** Representative images and quantification of red fluorescent protein (RFP) IHC on T128 metastasis-bearing livers. Area noted in yellow corresponds to T128 liver metastasis. **G, H** Quantification of surface liver metastasis of NCG mice intrasplenically injected with the T143 cell line that overexpresses RAB1A (n=4), EV (n=5), expresses RAB1A shRNA (n=8) or pLKO Control (n=8), Mann-Whitney test analysis. \**P* ≤0.05.

### Dual MNK/mTOR pharmacological inhibition decreases UM liver metastasis

To test the *in vivo* impact of combined MNKi-mTORi, T128 luciferase-tdTomato cells were intrasplenically injected into NCG mice (Fig. 5A). Drugs were administered by oral gavage 5 days post injection, to test the effect of the inhibitors on metastases already residing in the liver. Bi-weekly *in vivo* imaging of metastasis-bearing animals demonstrated a trend where treatment with SEL201 or INK128 alone decreased the luminescence of the liver compared to the vehicle treatment (Fig. 5B). However, only the combination treatment significantly impaired UM outgrowth in the liver, compared to the vehicle control group. The latter was confirmed at endpoint, where *ex vivo* imaging of livers demonstrated that the bioluminescence was decreased in the combination treatment, compared to single agents or vehicle cohorts (Fig. 5C). IHC staining of human nucleolin was performed to determine the percentage of human metastasis area within the livers, revealing that the combination treatment robustly decreased the metastasis area compared to the control (Fig. 5D). Furthermore, IHC staining revealed the on-target effects of SEL201 and INK128, showing supressed phosphorylation of eIF4E and S6, respectively (Fig. 5E, F). We had predicted that the combination treatment may function to decrease metastatic growth via decreased RAB1A expression, and we indeed found that RAB1A expression was repressed in metastases from the combination treatment arm, thereby supporting our hypothesis (Fig. 5G).

**Fig. 5.**
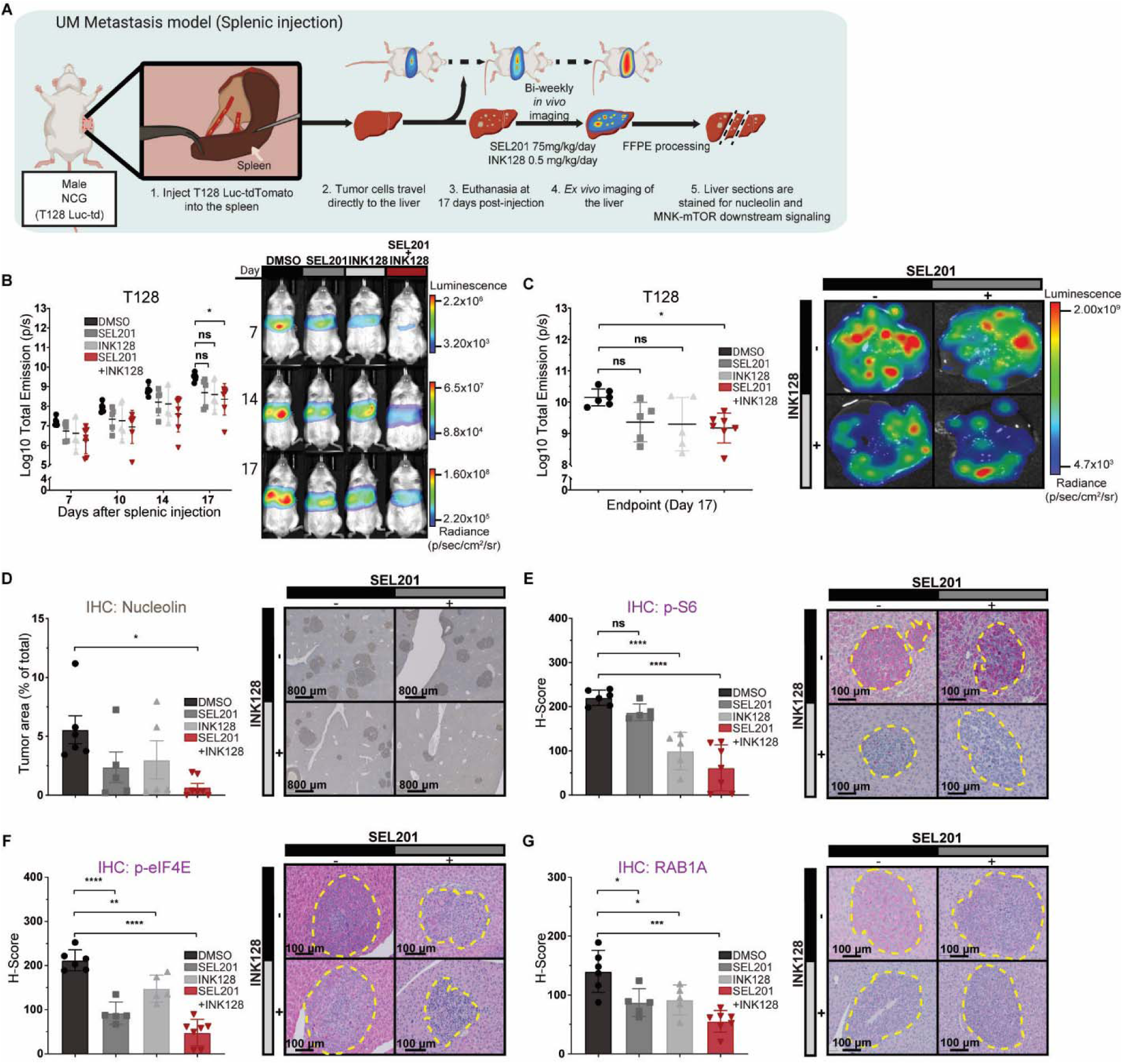
MNK1/2 and mTORC1/2 pharmacological inhibition reduces T128 liver metastasis *in vivo*. **A** Schematic overview of the splenic injection model using NCG mice injected with T128 luc-tdTomato; the tumor burden was determined by log10 luminoscore (photons/sec) acquired bi-weekly. NCG mice were treated with vehicle, SEL201 (75 mg/kg per day) and INK128 (0.5 mg/kg per day). **B** *In vivo* imaging of NCG bearing T128 luc-tdTomato tumors treated with vehicle (n=6), SEL201 (n=5), INK128 (n=5) and combination treatments (n=7), two-way ANOVA multiple comparisons test, ns=non-significant, \**P* ≤0.05 at day 17. **C** *Ex vivo* imaging of NCG with T128 luc-tdTomato metastases treated with vehicle (n=6), SEL201 (n=5), INK128 (n=5) and combination treatments (n=7), one-way ANOVA multiple comparisons test, ns=non-significant, \**P* ≤0.05. **D-G** Representative images and quantification of human nucleolin for metastasis area (detection using DAB), p-S6, p-eIF4E and RAB1A (detection using Magenta) IHC of NCG livers injected with T128 luc-tdTomato. One-way ANOVA multiple comparisons test, ns=non-significant, \**P* ≤0.05, \*\**P* ≤0.01, \*\*\**P* ≤0.001 and \*\*\*\**P* ≤0.0001. Areas noted in yellow correspond to T128 liver metastasis.

### The ER-to-Golgi secretory pathway is disrupted in RAB1A deficient UM cell lines

Evidence suggests that RAB1A increases the proliferation and invasion of cancer cells via multiple diverse mechanisms (56). Given that protein trafficking process was one of the pathways highlighted in our proteomic screen, and that RAB1A is known to coordinate ER-to-Golgi vesicle transport, we aimed next to determine whether the secretory pathway was disrupted in shRAB1A expressing UM cells. In eukaryotic cells, newly synthesized proteins transit through the secretory pathway, from the ER to Golgi stacks and finally to the trans-Golgi network (TGN). We employed the Retention Using Selective Hooks (RUSH) system, an assay that helps to synchronize protein secretion and visualize newly synthesized cargos through the secretory pathway (57). For the RUSH assay, we used the reporter cargo GPI-SBP-eGFP, which is retained in the ER due to the interaction between its streptavidin-binding peptide (SBP) and streptavidin in the hook protein (STR-Li:HA) we used (Fig. 6A). Upon the addition of biotin to the culture media, the biotin’s higher affinity for streptavidin competitively displaces the GPI-SBP-eGFP complex, allowing it to exit the ER and traffic across the secretory pathway. As expected, in the absence of biotin, we detected a similar colocalization of GPI-SBP-eGFP with the ER marker Hook (Li:HA) among the 3 cell lines (Fig. 6B, D). We next evaluated the colocalization of GPI-SBP-eGFP cargos with the cis-Golgi GM130 marker at 10, 20, and 30 min after the addition of biotin to monitor the cargo progression through the secretory pathway in pLKO control or shRAB1A T128 cells (Fig. 6C, E). We observed that cells with repressed RAB1A expression had a significant decrease in the co-localization of GM130 and GPI-SBP-eGFP cargos to the plasma membrane. These data suggest that upon repression of RAB1A there is impaired trafficking from the Golgi to the plasma membrane (Fig. 6E, F).

**Fig. 6.**
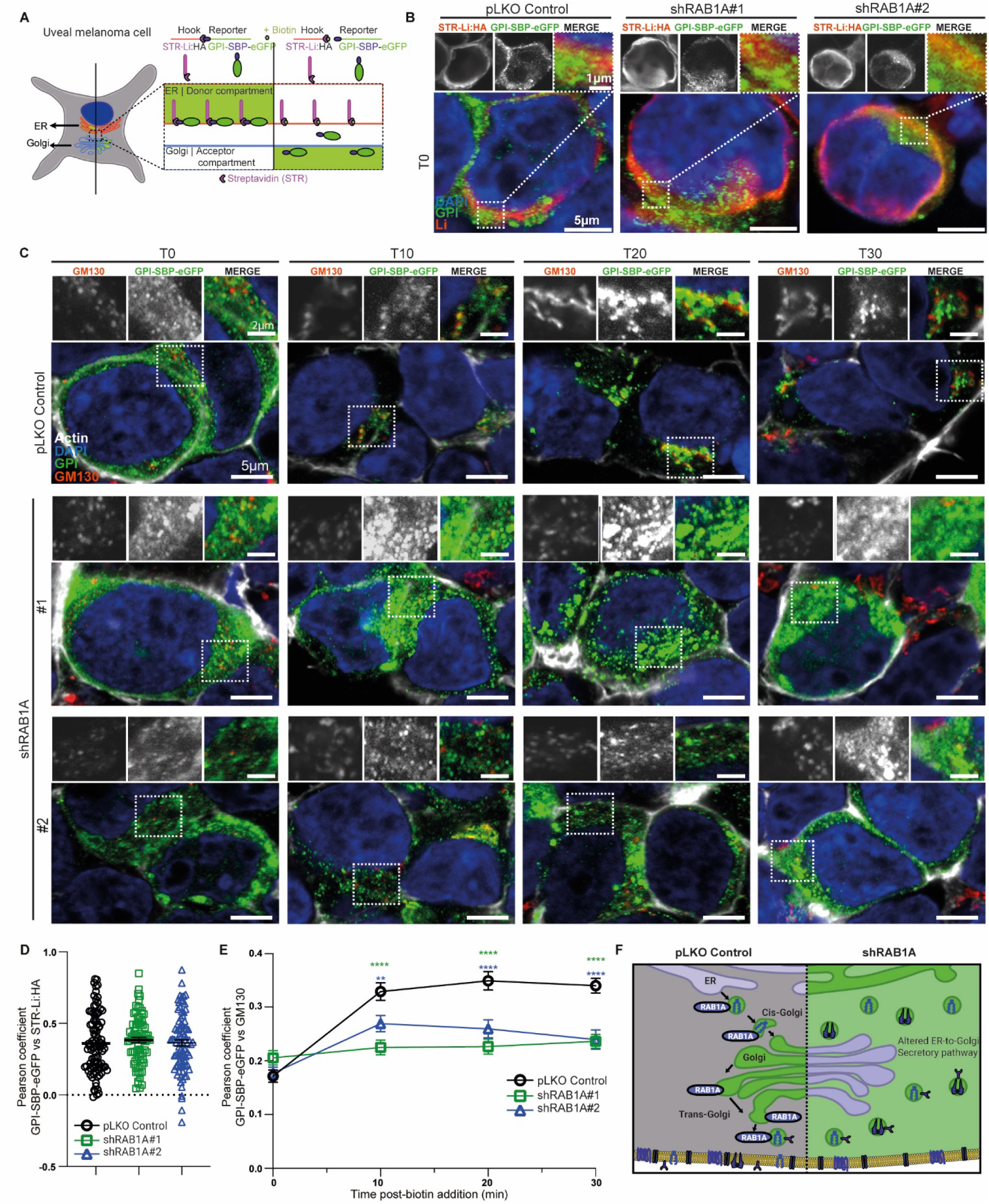
RAB1A depletion impairs the ER-to-Golgi secretory pathway. **A** Schematic overview of the RUSH assay imaging on T128 UM cell lines (pLKO and shRAB1A#1/2). Cells are transiently transfected with a plasmid containing the hook (STR-Li: HA) and the reporter (GPI-SBP-eGFP). Newly synthesized reporters are retained in the ER (donor compartment) by the hook, once biotin is added to the medium, reporters are released into the cytoplasm, allowing them to resume its trafficking to the Golgi. **B** Representative confocal images of cargo protein (GPI-SBP-eGFP, green) distribution and ER anchor protein (STR-Li:HA, red) prior to the addition of biotin (T0) in T128 cells. Single channels for Li and GPI are shown above the magnification of the boxed area displayed in the upper-right panel. **C** Representative confocal images of cis-Golgi (GM130, red) and cargo protein GPI-SBP-eGFP (green) at 0-, 10-, 20-, or 30-minutes post-addition of biotin to the culture medium. Individual and merged channels of the magnified boxed area are shown above each image. **D** Pearson coefficient correlation of STR-Li:HA and GPI-SBP-eGFP in c. **E** Pearson coefficient correlation of GM130 and GPI-SBP-eGFP at each timepoint of the RUSH assay. b, d: n=3 for pLKO Control, shRAB1A#1 n=3, shRAB1A#2 n=3. c, e: n=3. Error bars represent the mean ± S.E.M. One-way or two-way ANOVA with Dunnett post-comparison test was performed to compare RAB1A KD cells to the control. **F** Schematic describing the effect of decreased RAB1A expression on the secretory pathway, where it shows the improper localization of cargoes through the cytoplasm using the RUSH assay.

### Surfaceome profiling of UM cells with altered RAB1A expression uncovers novel druggable targets

Having determined that the RAB1A expression impacts the secretory pathway, we hypothesized that its expression may also alter plasma membrane-associated proteins. To test this, we examined the surfaceome, which is broadly defined as the subset of proteins expressed on the plasma membrane. We used a quantitative surface biotin-labeling approach coupled with mass spectrometry (Fig. 7A) and identified differentially expressed surface proteins in UM cells wherein RAB1A was either knocked down or overexpressed, compared to the respective controls (Fig. 7B and Supplementary Fig. S5A, B). We performed pathway analysis on the differentially regulated proteins and showed that RAB1A expression influences several signaling pathways previously shown to contribute to invasive and metastatic processes (Fig. 7C and Supplementary Fig. S5C-F). Among these, 12 proteins were downregulated upon RAB1A knockdown (shRAB1A) and conversely upregulated when RAB1A was overexpressed (RAB1A OE) (Fig. 7B, D). We next accessed the TCGA UMV dataset and showed that the identified surface proteins, cell adhesion molecule 1 (CADM1), ephrin type-B receptor 2 (EPHB2), and low-density lipoprotein receptor-related protein 8 (LRP8), positively correlated with RAB1A (Fig. 7E), uncovering novel candidate proteins for possible therapeutic intervention in UM. On the contrary, we noted a negative correlation of RAB1A with BAP1, a known tumor suppressor gene in UM. Supporting their potential clinical relevance in UM, we found that high expression of CADM1, EPHB2, and LRP8 were significantly associated with poorer overall survival in patients (Fig. 7F-H). These findings suggest that RAB1A abundance plays a role in shaping the UM surfaceome, providing potential druggable targets in UM.

**Fig. 7.**
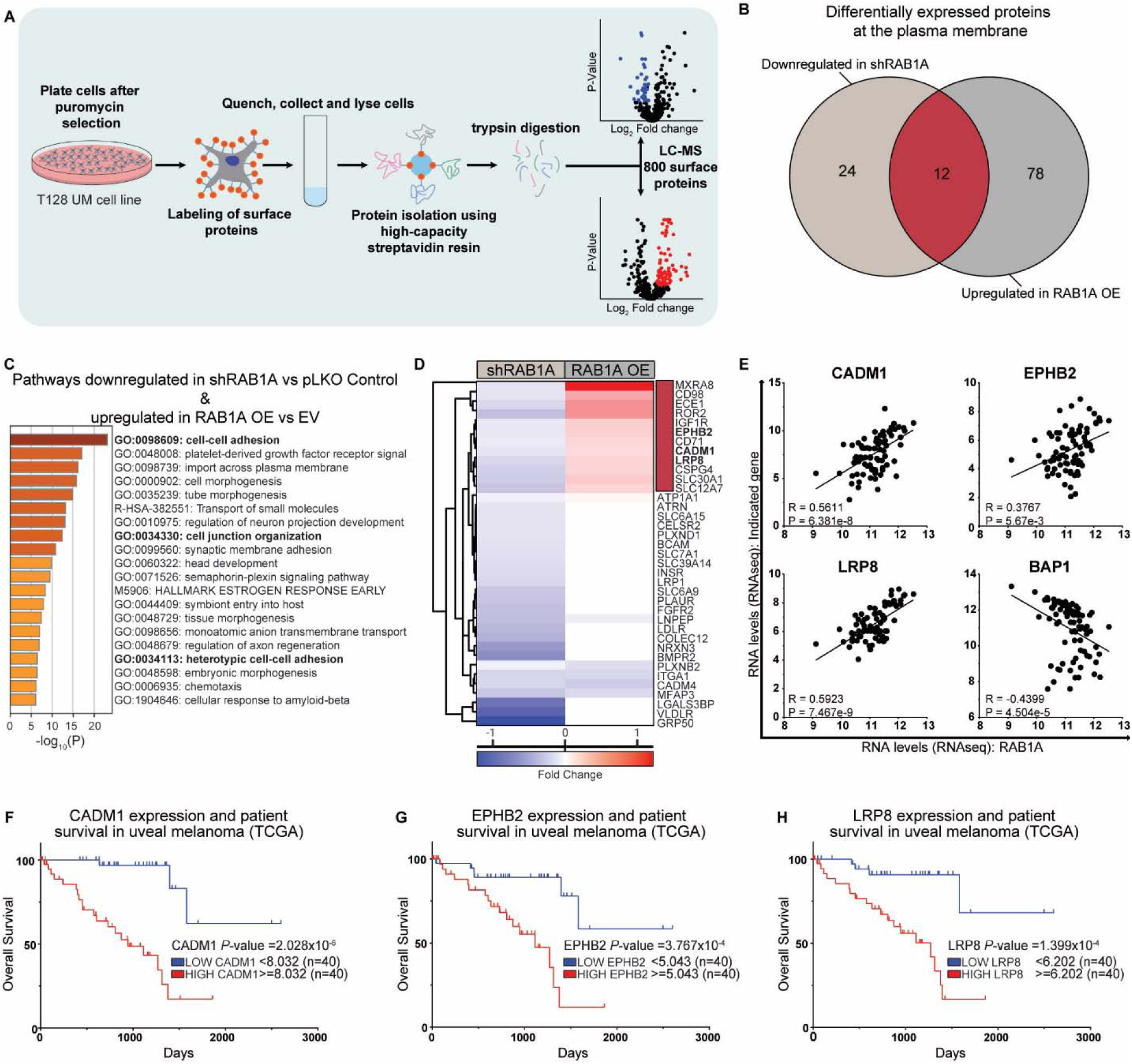
Surfaceome profiling of UM cells with altered RAB1A expression uncovers novel druggable targets. **A** Schematic of the methodology followed to determine protein expression at the cell surface in T128 RAB1A OE, EV, shRAB1A and pLKO Control UM cell lines. LC-MS: Liquid chromatography mass spectrometry. **B** Venn diagram describing the number of differentially expressed proteins at the cell membrane between the indicated cell lines. **C** Pathway enrichment analysis of proteins at the cell membrane found to be downregulated in **B** shRAB1A vs pLKO Control and upregulated in RAB1A OE vs EV. **D** Heatmap of hierarchical clustering of proteins downregulated in shRAB1A and upregulated in RAB1A OE at the cell membrane. **E** Correlation of RAB1A RNA expression with the RNA expression of the listed genes. Spearman-rank order, linear regression shown. **F-H** Kaplan-Meier overall survival plots depicting CADM1, EPHB2, LRP8 and BAP1 expression in the TCGA UVM dataset (n=80).

## DISCUSSION

The assembly of the eIF4F complex has been shown to be druggable in other tumor contexts and is also the converging point of two major signalling pathways in UM − MAPK and PI3K (31, 44). Our data show that both axes cooperate to enhance metastatic potential and, when inhibited simultaneously, the result is repression of clonogenicity and invasivity *in vitro*, and decreased experimental UM liver metastasis *in vivo*.

In this study, we identify RAB1A as a critical downstream effector of activated MNK and mTOR. Other studies have highlighted diverse mechanisms of action associated with the anti-cancer benefits of combined MNK+mTOR inhibition. Teo *et al*. showed that suppression of MNK2 (MKI4) and MNK1/2 (MNKI57) with mTOR(rapamycin) resulted in decreased proliferation in leukemic cells, but only observed G1 cell cycle arrest when inhibiting MNK1/2 simultaneously with rapamycin (27). A similar phenotype was also observed in prostate cancer models, where the inhibition of MNK1/2 (CGP57380) and mTOR together induced G_1_ cell cycle arrest due to decreased expression of cyclins A, B, and D1 (13). A recent study by Knight *et al.*, performed in *KRAS*-mutated colorectal cancer, showed robust *in vivo* anti-tumor benefit of combined MNK (eFT508)-mTOR (rapamycin) inhibition, where reducing protein synthesis and blocking p-eIF4E resulted in repressed c-MYC expression (25). While both our study and Knight *et al*. focused on the tumor-intrinsic effects of MNK1/2 inhibition; numerous studies are emerging that point us to appreciate the role of MNK-eIF4E axis in cells of the tumor microenvironment (TME) (58). It was shown recently that the MNK-eIF4E axis cooperates with mTOR to shape prostate cancer translatome, producing soluble factors (BGN, SPP1, and HGF) that attract myeloid-derived suppressor cells (MDSCs) (26). Unfortunately, the lack of an immune competent mouse model for UM hinders our ability to characterize the impact of blocking MNK and mTOR on cells of the TME. We anticipate that with such a pre-clinical mouse model in hand, we will be able to expand upon prior work showing that blocking the MNK-eIF4E axis can potentiate host anti-tumor immunity. For example, the MNK-eIF4E axis can regulate the expression of PD-L1 on cancer cells(59), dendritic cells, and MDSCs (30). We eagerly await the development of an immune competent model of metastatic UM.

Proper ER to Golgi-mediated protein vesicle trafficking is essential to maintain cell homeostasis, controlling tightly regulated processes such as protein location to the organelles, molecule release to the extracellular compartment (60), internalizing cell membrane receptors and protein degradation via lysosomes (61). Nevertheless, it can be hijacked by cancer cells to impact the steps in the metastatic cascade such as modifying the tumor microenvironment, aberrant pathway signalization and enhancing oncogenic activity (61, 62). RABs control the major steps of vesicle trafficking by switching from a GDP-inactive to a GTP-active state (mediated by diverse guanine exchange factors [GEFs]) (63). Once GTP bound, RABs will recruit effectors to regulate most steps of vesicular transport, from cargo selection to vesicle budding, transport, tethering and fusion (62, 64). Deregulated cargo delivery and altered RAB family expression can be exploited by cancer cells, where dysregulated vesicle localization triggers metastatic growth (65). One example is RAB40b, as its localization in secretory vesicles has been shown to be critical for the sorting and transport of MMP2/9. When RAB40b expression is altered, MMPs are directed to lysosomes, resulting in a decreased potential to degrade the extracellular matrix in *in vitro* breast cancer models (66). Another mechanism by which RABs are hijacked in cancer is the overexpression of RAB25 in radio-resistant lung cancer. The interaction of RAB25 with the EGFR decreased its degradation and enhanced the recycling of the receptor to cell surface in radioresistant lung adenocarcinoma (LUAD) (67). There are many examples of how RABs have a role in metastatic disease. Although the development of selective inhibitors targeting specific RABs is difficult to achieve due to the close homology among all RAB proteins, the team of Shokat *et al.* recently sought to optimize inhibitors of RAB1A, which may hold promise in UM (68). Here, we describe that increased RAB1A expression correlates with p-eIF4E and p-S6 in UM and results in decreased patient survival. Moreover, using combined pharmacologic inhibition of MNK and mTOR, we blocked RAB1A expression in *in vitro* and *in vivo* models of UM. Our research, and others, indicate that RAB1A seems to play a crucial role in promoting tumorigenesis, where its overexpression can be used as prognostic marker in colorectal and lung (NSCLC) cancers (51, 56, 69). Mechanistically, we showed that when RAB1A expression is decreased, there is a defect in the transport of GPI-SBP-eGFP cargos through the secretory pathway. In our context, we propose that this could lead to altered expression of cell surface proteins, potentially influencing the malignant behavior of cancer cells. Many studies have highlighted the importance of understanding cell surface dynamics in disease progression, particularly how oncogenic kinases such as AKT and MEK drive functional changes that impact invasive and adhesive processes. Some of these alterations can be reversed upon MEK inhibition, highlighting the role of MAPK signaling in surfaceome remodeling (70, 71). Here, using UM cells with genetically manipulated RAB1A expression, we demonstrated that RAB1A abundance influences the expression of CADM1, EPHB2, and LRP8. We then used publicly available data to show a positive correlation of CADM1, EPHB2, and LRP8 with RAB1A, and finally that the expression of those identified plasma membrane-associated proteins was correlated with worse overall survival in UM. Our findings further support previous observations that RAB1A regulates the abundance and location of surface markers at the plasma membrane. Notably, in other contexts, RAB1A influenced migration through integrin β1 recycling and localization to lipid rafts (72). In a similar fashion, our results highlight possible novel proteins that may have clinical relevance, suggesting that targeting RAB1A could offer novel therapeutic opportunities in UM.

## ACKNOWLEDGMENTS

Procurement of UM cells was possible thanks to the common infrastructure “Uveal Melanoma Biobank”, which is financially supported by the Fonds de recherche du Québec - Santé (FRQS)-Vision Sciences Research Network (VSRN; SL). SL is a Junior 2 Research Scholar of the FRQS. The authors wish to extend their gratitude to Mr. Christian Young, manager of the Flow Cytometry Core facility at the Lady Davis Institute and Dr. Naciba Benlimame of the Molecular Pathology Core of the JGH, for their technical support. We express our appreciation to Mrs. Véronique Michaud and Mrs. Kathy Ann Forner for their aid into developing our animal models. We acknowledge the technical support of the Lady Davis Institute for Medical Research Imaging and Phenotyping Core. We are grateful for the support from Mr. Marc Bazin, the platform manager of the In Vivo Imaging System Lumina (Centre de recherche du CHU de Québec-Université Laval), for its collaboration to acquire luminescent images

## ETHICAL APPROVAL

All animal experiments described were carried out in compliance with the Canadian Council of Animal Care guidelines and regulations. Animal protocols were reviewed and approved by McGill University Animal Care and Use Committee (JGH-8168)

