## Supplemental figures and tables for "RAB1A is a novel vulnerability in uveal melanoma revealed by dual inhibition of MNK1/2 and mTOR"

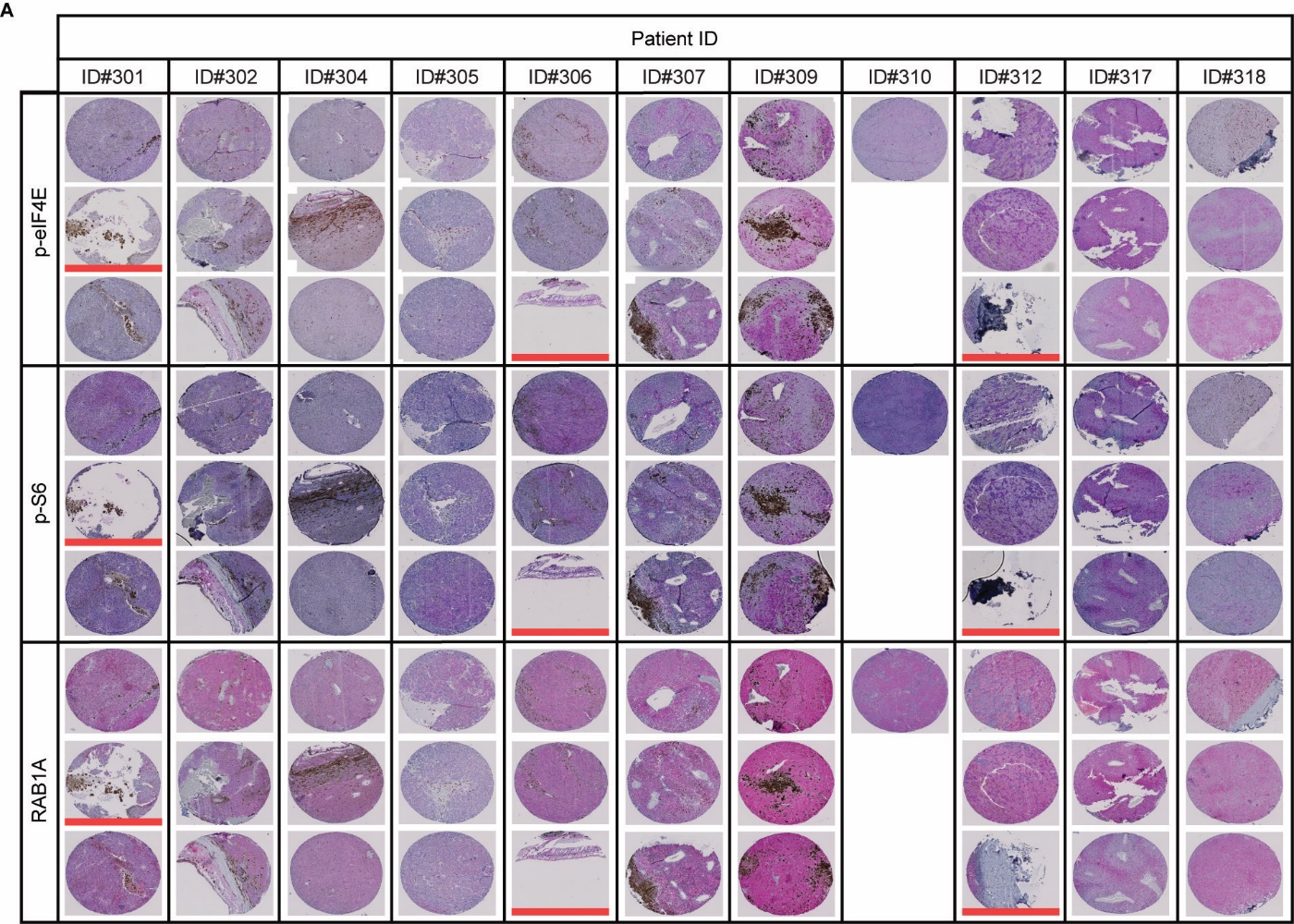
**Supplementary Fig. S1 MNK1/2-mTORC1/2 activation status in primary-derived human UM TMA.** **A** IHC stains of p-eIF4E and p-S6 on primary human UM TMA (n=28 cores). The cores were stained and graphed using pixel intensity and organized using patient ID, and later scored using 3 different level of expression, corresponding to low (negative control stain), medium or high expressing. Exhausted cores shown with red underline were omitted from the analysis (n=3 cores).

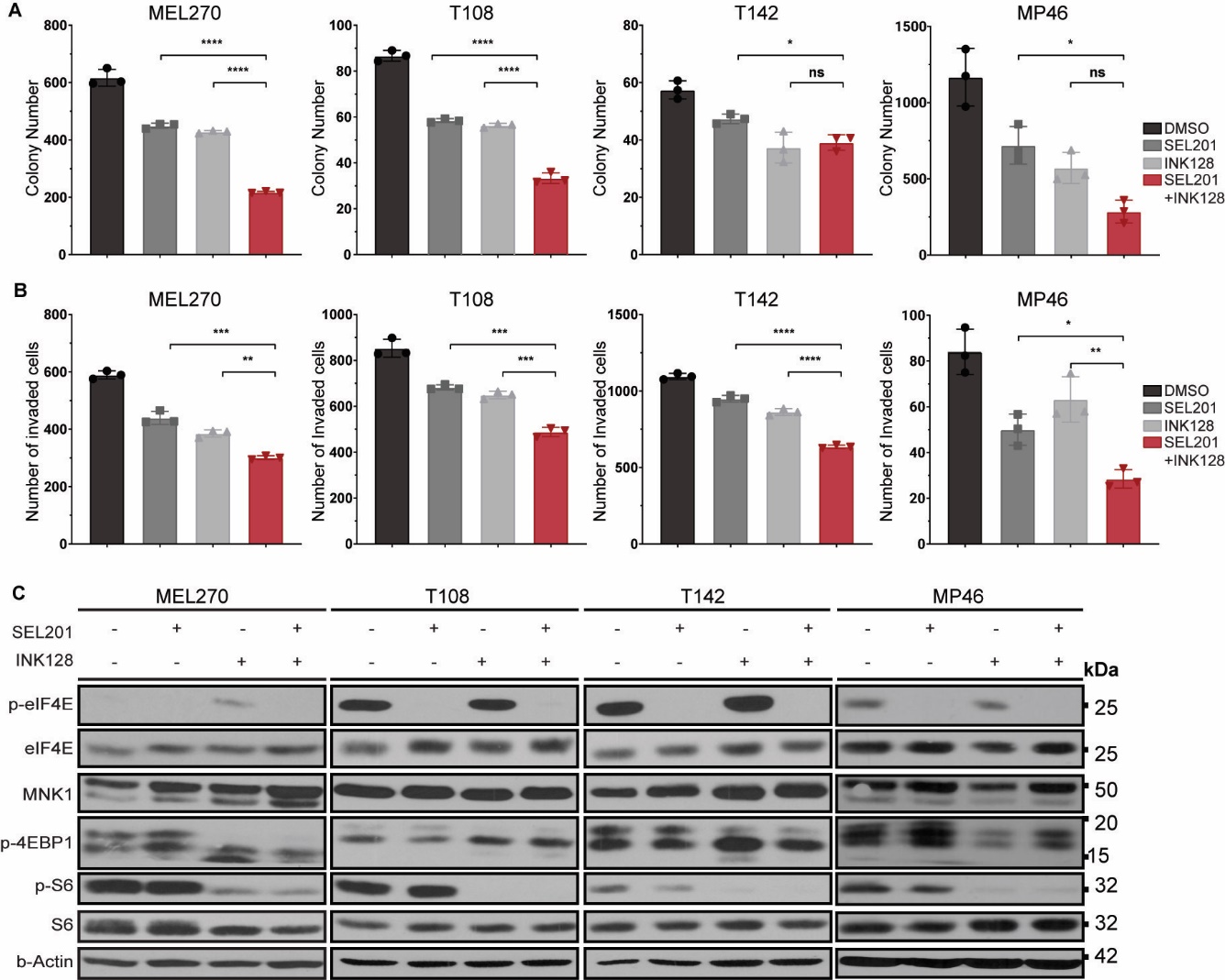
**Supplementary Fig. S2** **MNK1-2 and mTORC1/2 pharmacological inhibition decreases UM clonogenic and invasive outgrowth *in vitro*.** **A, B** The inhibitor SEL201 cooperates with INK128 to decrease clonogenicity and invasion outgrowth in UM cell lines. Experiments represent the mean ±SD of 3 experimental replicates performed in technical triplicate. One-way ANOVA multiple comparisons test. **P* ≤0.05, ***P* ≤0.0, ****P* ≤0.001 and *****P* ≤0.0001 compared to single treatments. **C** Immunoblot analysis of the indicated proteins in MEL270, T108, T142 and MP46 UM cell lines treated with SEL201 and INK128 inhibitors, alone or in combination.

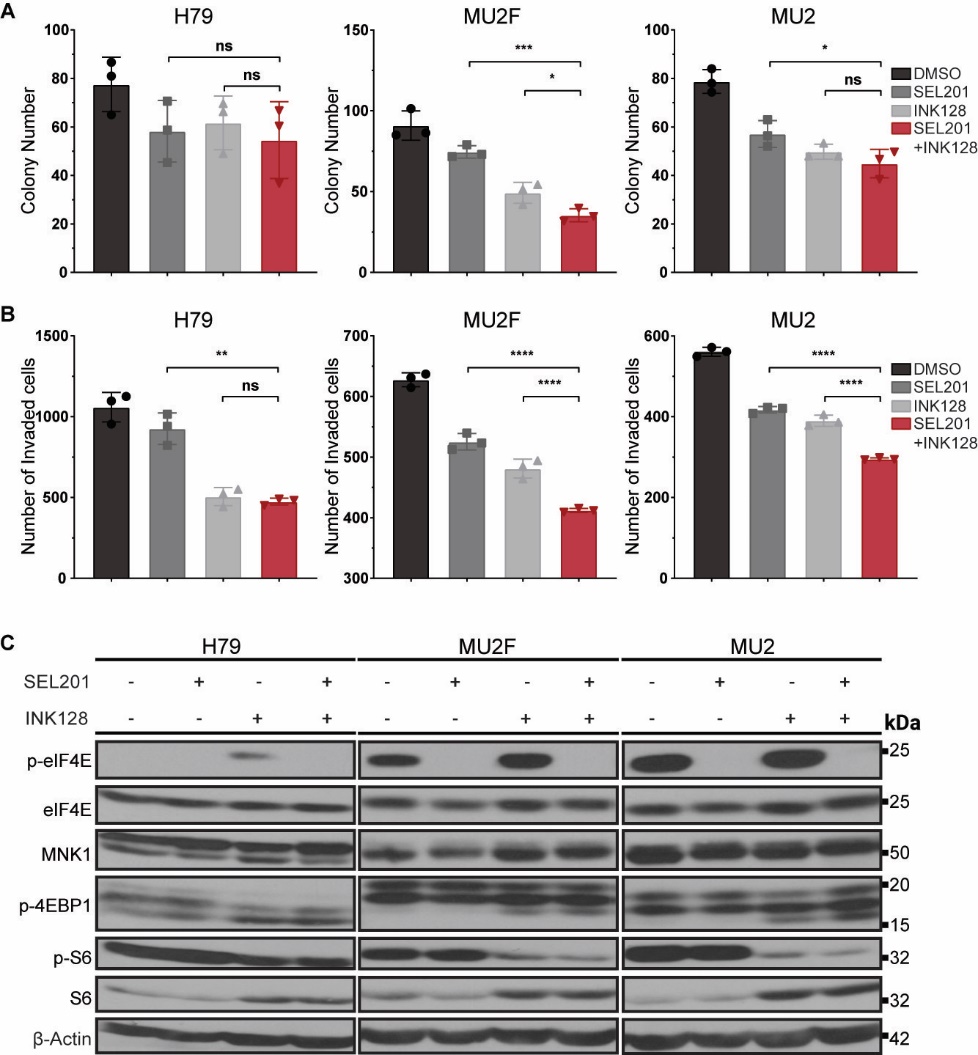
**Supplementary Fig. S3 MNK1-2 and mTORC1/2 pharmacological inhibition decreases UM clonogenic and invasive outgrowth *in vitro*.** **A, B** The inhibitor SEL201 cooperates with INK128 to decrease clonogenicity and invasion outgrowth in UM cell lines. Experiments represent the mean ±SD of 3 experimental replicates performed in technical triplicate. One-way ANOVA multiple comparisons test. **P* ≤0.05, ***P* ≤0.01, ****P* ≤0.001 and *****P* ≤0.0001 compared to single treatments. **C** Immunoblot analysis of the indicated proteins in H79, MU2F and MU2 UM cell lines using SEL201 and INK128 inhibitors, alone or in combination.

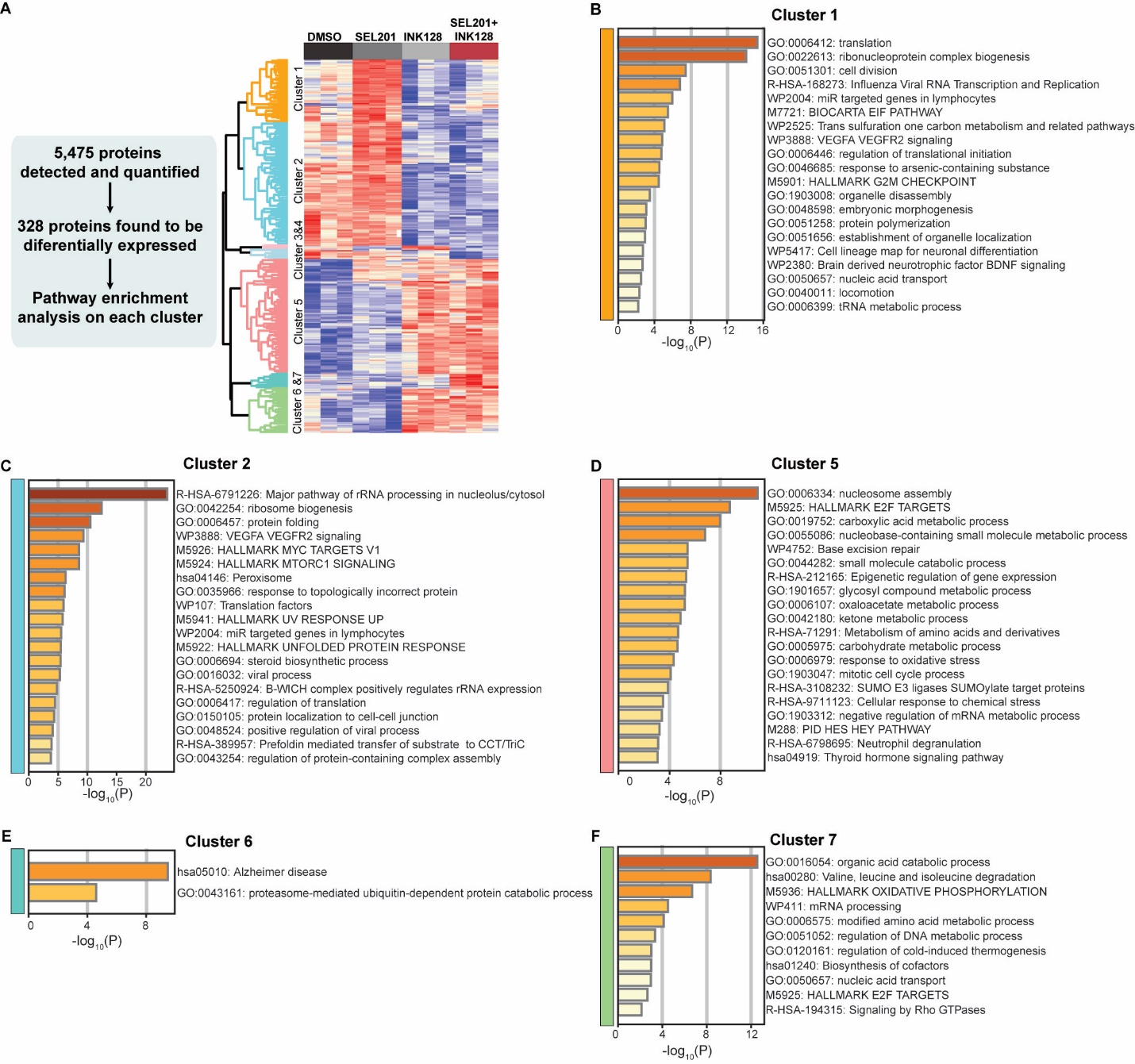
**Supplementary Fig. S4** **Proteome profiling of T128 UM cell line shows that inhibition of MNK1/2 and mTORC1/2, alone or in combination, impacts multiple signaling pathways.** **A** Schematic of downstream analysis and hierarchical clustering of the differentially regulated proteins detected using DIA on UM cells treated with vehicle, SEL201, INK128 or combination therapy at 24 hours. **B** Pathway enrichment analysis of cluster 1. **C** Pathways enrichment analysis of cluster 2. **D** Pathway enrichment analysis of cluster 5. **E** Pathway enrichment analysis of cluster 6. **F** Pathway enrichment analysis of cluster 7. Pathway enrichment analysis was not performed on cluster 3 due to small proteins number.

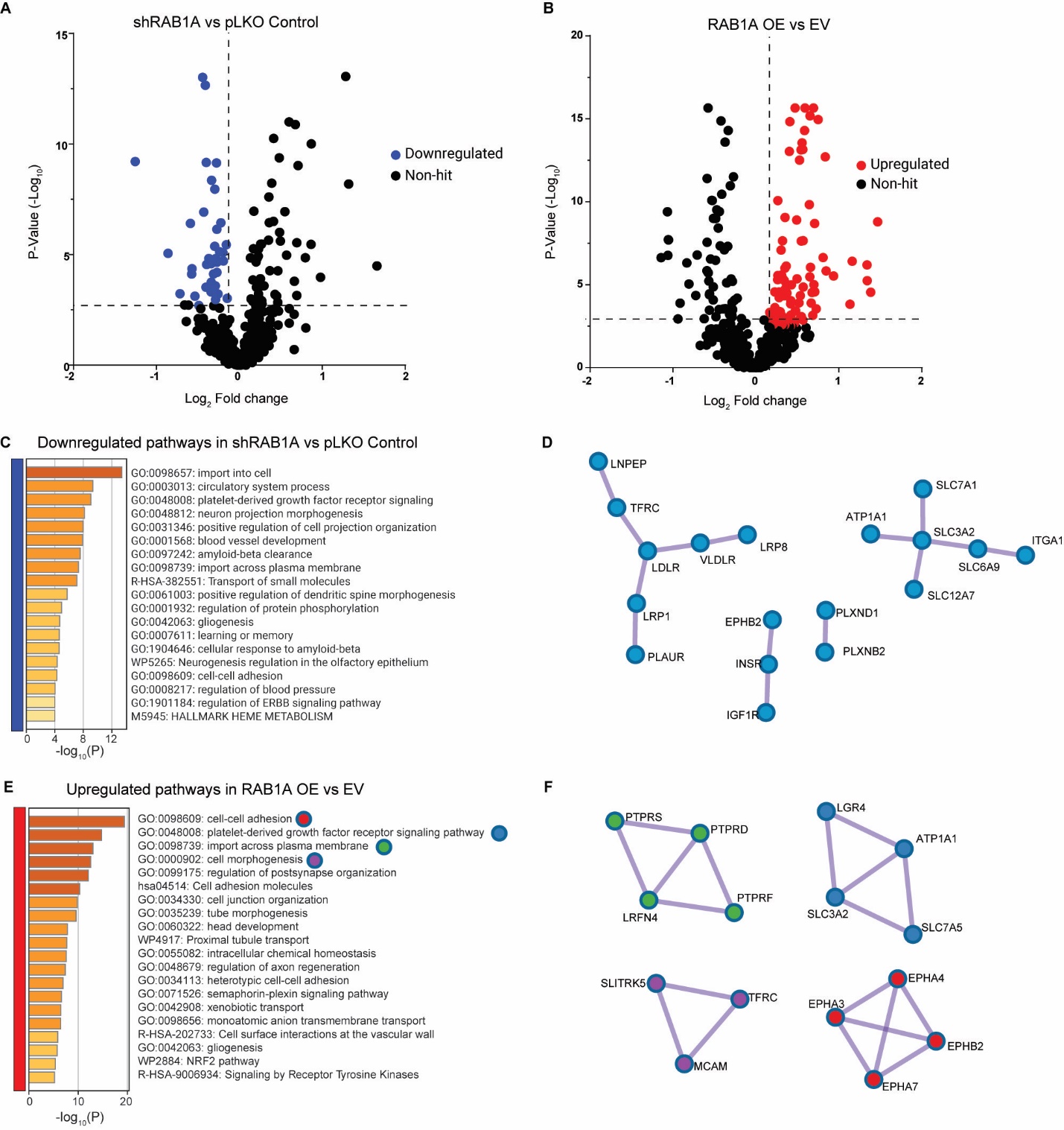
**Supplementary Fig. S5** **Surfaceome analysis of differentially regulated proteins in shRAB1A and RAB1A OE vs control.** **A, B** Volcano plots showing the differential protein expression at the plasma membrane of T128 cells transduced with shRNA against RAB1A compared to pLKO Control and RAB1A OE compared to empty vector (EV). Dashed lines depict significance (*P* < 0.001) and fold enrichment ≤+ and - 0.15 thresholds. **C, D** Pathway enrichment analysis and protein-protein interaction of surface markers downregulated in shRAB1A compared to pLKO Control. **E,** **F** Pathway enrichment analysis and protein-protein interaction of surface markers upregulated in RAB1A OE compared to EV.to EV.

SUPPLEMENTAL TABLES

Supplemental table 1. Information of primary antibodies used for immunoblotting, immunohistochemistry and immunofluorescence assays.

| **Target** | **Antibody Source and Catalog (#)** | **Application** | **Dilution** |
| --- | --- | --- | --- |
| p-eIF4E | Cell Signaling #9741 | Immunoblot | 1:1000 |
| eIF4E | BD Biosciences #610270 | Immunoblot | 1:1000 |
| MNK1 | Cell Signaling #2195 | Immunoblot | 1:1000 |
| p-4EBP1 | Cell Signaling #2855 | Immunoblot | 1:1000 |
| 4EBP1 | Cell Signaling #9452 | Immunoblot | 1:1000 |
| p-S6 | Cell Signaling #5364 | Immunoblot | 1:1000 |
| S6 | Cell Signaling #2317 | Immunoblot | 1:1000 |
| β-Actin | Sigma Aldrich #A5441 | Immunoblot | 1:5000 |
| RAB1A | Proteintech #11671-1-AP | Immunoblot | 1:1000 |
| p-eIF4E | Abcam #ab76256 | Immunohistochemistry | 1:50 |
| p-S6 | Cell Signaling #35708 | Immunohistochemistry | 1:200 |
| RAB1A | Abcam # ab302545 | Immunohistochemistry | 1:100 |
| Nucleolin | Abcam # ab136649 | Immunohistochemistry | 1:2000 |
| RFP | Rockland #600-401-379 | Immunohistochemistry | 1:300 |
| HA-Tag | Cell Signaling #3724 | Immunofluorescence | 1:800 |
| Anti-GFP | Santa Cruz #sc9996 | Immunofluorescence | 1:1000 |
| Anti-GM130 | Cell Signaling #12480 | Immunofluorescence | 1:3000 |
